# A duplexed CRISPRi-based approach to predict synergistic antibiotic targets in *Escherichia coli*

**DOI:** 10.1101/2024.05.15.594409

**Authors:** Xianyi Xiong, Hannah Furseth, Brady McCoy, Lauren Slavic, Xun Yuan, Sydney Chen, Anthony D. Baughn, Yusuke Minato, William R. Harcombe

## Abstract

Co-repressing drug targets with CRISPR interference (CRISPRi) has the potential to predict both synergistic and additive drug combinations. In a screen experiment with *Escherichia coli*, pairwise gene co-repression assays correctly predicted the interaction type observed for 36 antibiotic pairs in checkerboard assays. Furthermore, we used this approach to identify promising synergistic interactions with a gene currently not targeted by clinical antibiotics. CRISPRi can systematically test the impact of gene inhibitions and identify novel drug targets.

## Main Text

Synergistic drug combinations have higher efficacy than predicted from each drug’s individual effects (1). Used clinically, these drug synergies can increase effectiveness at reduced drug dosages and alleviate side effects (1,2). Despite these benefits, it remains challenging to predict novel synergistic gene target pairs to better treat infections (3).

Drug target identification and obtaining new synergistic drug pairs often rely on chemical genomics (2,4), which has many limitations. Chemical genomic assays frequently involve treating a genome-wide, loss-of-function mutant library of a pathogen with a known drug at a sublethal concentration, and measuring fitness for each mutant in the library (4). Chemical genomic assays can help study the genetics of altered antibiotic susceptibility (4-6), infer mechanisms for antibiotic synergies (5,7,8), and predict potential synergistic target pairs for future validation (9-11). However, the role of essential genes cannot be studied using loss-of-function mutant libraries because cells without essential functions cannot survive and therefore are missing from the libraries (12). Second, effects from loss-of-function mutations in genes may be distinct from antibiotic activity as antibiotics can be titrated by varying their concentrations (2). So it is unclear whether the gene-to-drug interactions measured by altered drug susceptibilities for gene knockout mutants is a similar enough measurement to the drug-to-drug interactions we intend to understand.

CRISPR interference (CRISPRi) can expand chemical genomics assays for studying how slightly perturbing an essential gene’s expression affects drug susceptibility (13-16). CRISPRi uses a small guide RNA (sgRNA) that directs a deactivated Cas9 (dCas9) endonuclease to a gene target through sequence complementarity, and represses transcription of the target gene without creating a double-stranded break (17). Unfortunately, chemical genomic studies using CRISPRi libraries are still constrained to identifying synergistic targets to existing drugs and neglect truly novel target pairs in the genome (3). However, because multiple sgRNA sequences can be expressed together to target more than one gene (multiplexed CRISPRi), epistatic interactions from perturbing multiple genes simultaneously can be measured. Thus, we hypothesize that duplexed CRISPRi–whereby two target genes are simultaneously repressed–can be directly used to predict interactions between drugs targeting the product of each gene.

It was previously shown that duplexed CRISPRi can help identify new targets to potentiate existing antibiotics (18-20). However, it remains challenging to distinguish synergistic from additive (i.e. non-potentiating) drug interactions (21). Here, we showed that duplexed CRISPRi can directly distinguish between synergistic and additive interactions in *Escherichia coli*. We then used duplexed CRISPRi to predict an under-characterized gene target that can potentially synergize with others in the folate biosynthesis pathway.

### Duplexed CRISPRi reliably predicts synergistic combinations in antibiotic pairs

*E. coli* LC-E75 is an MG1655 derivative with a dCas9 gene that is encoded in the genome and inducible with anhydrotetracycline (ATc) (22). To simultaneously repress transcription for two genes and measure the epistatic interaction of co-repression, we constructed a plasmid with two vector sgRNA sequences that can be modified separately via Golden Gate Assembly (23) (Fig. 1A and S1A-B, Supplementary Methods 1-2). We validated that targeting the same essential gene at either sgRNA site is equally effective (Fig. S1C). To mimic sub-lethal effects of antibiotics, we chose sgRNAs that lead to weak transcriptional inhibitions in LC-E75 in liquid LB medium as previously described (22) (Supplementary Method 3). The fitness effect of transcriptional repression in each strain was measured as relative yield (CFU) induced by 10 nM ATc after 24-hour growth in serially diluted spots plated on M9 agar supplemented with glucose (0.2% m/v). The interaction index between each pairwise gene knockdowns (*S*) was calculated using a Bliss independence metric (Fig. 1A and S1D) (24). We considered an interaction synergistic (antagonistic) if knocking down the same two genes consistently leads to high (low) average *S* values, 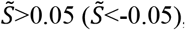, over three biological replicates (Fig. 1A). Additivity is claimed if the interaction is neither synergistic nor antagonistic.

**FIG 1.**
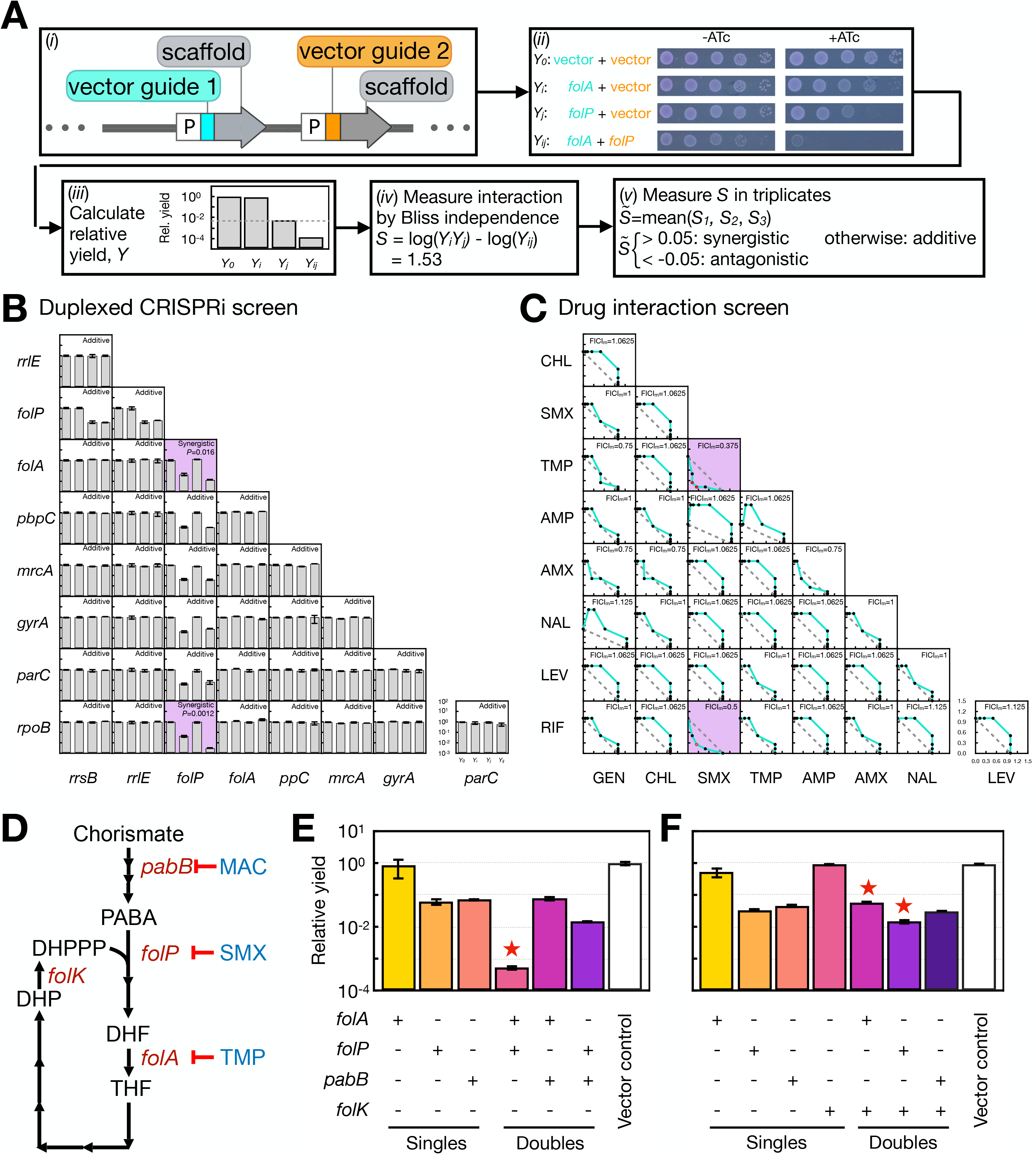
Duplexed CRISPRi can distinguish between synergistic target pairs from additive ones in *E. coli*. (A) Schematic for measuring gene-to-gene interactions from duplexed CRISPRi knockdowns. Step (*i*): Reducing transcription using duplexed CRISPRi by targeting one or two genes with sgRNAs on a plasmid. P: a constitutive promoter. Step (*ii*): Fitness effect caused by transcriptional repression of gene targets was measured on M9-glucose agar with and without dCas9 induction by 10 nM ATc. Fitness was measured for strains with no gene knock down (*Y*_*0*_), with a single gene knocked down (*Y*_*i*_ or *Y*_*j*_), and with both genes knocked down (*Y*_*ij*_). Vector: vector sgRNA; *folA, folP*: example gene targets. Step (*iii*): Fitness defect was measured as CFU yield reduction due to dCas9 induction. Grey line: expected fitness of knocking down both genes calculated as *Y*_*i*_*Y*_*j*_. Step (*iv*): The interaction between a transcriptional repression pair was measured by a log-transformed Bliss independence metric, *S*. Step (*v*): Synergy is claimed (one-sided t-test, *P*<0.05, same throughput manuscript) by comparing mean of three independent *S* measurements 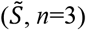 to a threshold 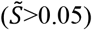. Antagonism is claimed if 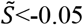 is statistically supported. Additivity is claimed if the interaction is neither synergistic nor antagonistic. (B) Epistatic interaction type for transcriptional co-repressions by duplexed CRISPRi in 36 pairwise gene combinations were measured as in Fig. 1A. Each gene corresponds to one antibiotic target in Fig. 1C. Clear and non-clear backgrounds indicate additive and synergistic interactions, respectively. *Y*-axis in each plot denotes the relative yield due to ATc induction. Other than the *folA-folP* pair (*P*=0.016), the *folP-rpoB* pair (*P*=0.0012) also appears synergistic in the CRISPRi screen. (C) Pairwise antibiotic interactions for nine antibiotics were measured with the fractional inhibitory concentration indices (FICI) for each drug combination in a checkerboard assay. Each axis denotes the fractional inhibitory concentration (FIC) an antibiotic. Drug combinations with small minimum FICI values (FICI_m_≤0.5) are considered synergistic pairs (non-clear background), while FICI_m_>4 indicate antagonism, and 0.5<FICI_m_≤4 are considered additive/indifference (8,21). Red dots: antibiotic concentration combinations with FICI≤0.5. Fig. S2 contains full names, mechanisms, gene targets, and MICs in M9-glucose media for all nine antibiotics used. An interval on each axis denotes 0.3 FIC. (D) Schematic of inhibitors targeting the folate biosynthesis pathway in *E. coli* as described previously (8). (E-F) Co-repressing the *folA-folP* pair results in synergistic effect (*P*=0.0092), but not for the *folA-pabB* (*P*=0.80) or *folP-pabB* (*P*=0.99) pairs. Repressing the *folA-folK* (*P*=0.0016) and *folP-folK* (*P*=0.025) pairs results in synergistic effects, but repressing the *pabB-folK* pair does not (*P*=0.19). “+” signs below each bar indicates transcriptional repression. Red star indicates synergistic repression pairs.

We first demonstrated that duplexed CRISPRi can accurately predict drug interactions across broad antibiotic classes. For nine antibiotics spanning five major clinically relevant classes (25), we identified one major gene target for each and constructed plasmids to repress transcription of each gene individually or in pairs (Fig. S2A). Duplexed CRISPRi predicted that among all 36 gene repression pairs, the only two synergistic combinations in *E. coli* LC-E75 are *folA-folP* and *folP*-*rpoB*, whereas all remainder interactions are additive (Fig. 1B; Supplementary Data). These predictions were confirmed by measuring fractional inhibitory concentration indices (FICI) in LC-E75 using checkerboard assays (Fig. S2) (26). The sulfamethoxazole-trimethoprim (SMX-TMP) and sulfamethoxazole-rifampin (SMX-RIF) antibiotic pairs are synergistic and borderline synergistic, respectively (Fig. 1C). No false-positive hit was found as all other drug interactions were classified as additive.

### Using duplexed CRISPRi to explore novel synergistic target pairs

We used duplexed CRISPRi to search for underexplored synergistic gene target pairs. Synergistic drug target pairs are hypothesized to be common in metabolic loops wherein downstream metabolic products are also substrates used to synthesize upstream molecules (8). For example, all three antibiotics of SMX (*folP* inhibitor), TMP (*folA* inhibitor), and MAC (MAC173979, *pabB* inhibitor) inhibit steps in the folate biosynthesis pathway (Fig. 1D) (8,27). However, SMX-TMP is the only synergistic pair among the three antibiotics, because both drugs can reduce the flux through the steps they target and hence potentiate the other drug’s activity—a mechanism contributing to synergy called “mutual potentiation” (8). We found that repressing the gene pair *folA*-*folP* using duplexed CRISPRi also leads to a strong synergistic effect, whereas knocking down the *folA-pabB* or *folP-pabB* pairs does not (Fig. 1E). Polar effects are a concern for CRISPRi assays, where repressing transcription of genes early in an operon will also affect genes that lie downstream (17). Here polar effects are unlikely to explain the *folA-folP* synergy, because repressing *glmM*, the gene after *folP* in the operon, does not lead to a synergistic effect with *folA* (Fig. S3).

To test whether other synergistic gene target pairs exist within the same metabolic loop in folate biosynthesis (8), we co-repressed the transcription of a non-essential gene, *folK* (28), with the three others explored above (Fig. 1D). We found synergistic interactions for the *folA-folK* and *folP-folK* pairs but not for the *pabB-folK* pair (Fig. 1F). This evidence implies that mutual potentiation is a more common mechanism contributing to drug synergy in bacteria than previously thought. Future work should acquire *folK* inhibitors via chemical synthesis or from chemical libraries to consolidate these predictions.

In this study, we validate that duplexed CRISPRi can distinguish between synergistic and additive drug interactions across many antibiotic classes in *E. coli*. This method can potentially be scaled up to systematically survey gene-to-gene epistatic interactions and holistic prediction of synergistic target pairs. While polar effects are a concern for genome-wide CRISPRi screen assays (13,17,22), they can be further avoided by performing duplexed CRISPRi to investigate epistatic operon-to-operon interactions, which will still result in a rich dataset of >240,000 pairwise combinations from the estimated 600∼700 operons in the *E. coli* genome (29).

## Acknowledgements

We thank A.T. Bisesi, J.N.V. Martinson, J.M. Chacón, A. Narayanan (U of MN), M. Howe (Mayo Clinic), J.M. Rock (Rockefeller) for the helpful feedback. We thank D. Bikard for gifting us the *Escherichia coli* LC-E75 strain (Addgene #115925) and the psgRNA plasmid (Addgene #114005).

## Author Contributions

X.X. conceived the study, designed the study, performed data analysis, and wrote and edited the manuscript. H.F., B.M., and L.E.S. designed the study, performed data collection and analysis, and contributed to manuscript editing. X.Y. and S.C. contributed significantly to data collection. A.D.B. and Y.M. partly supervised the study and edited the manuscript. W.R.H. supervised the whole study, acquired and managed funding, and edited the manuscript.

